# Modulating cerebrospinal fluid dynamics using pulsed photobiomodulation: feasibility, parameter and skin-colour dependence

**DOI:** 10.1101/2025.05.06.652458

**Authors:** Ariel Motsenyat, Xiaole Z. Zhong, Hannah Van Lankveld, Joanna X. Chen, Alicia Mathew, J. Jean Chen

## Abstract

The use of photobiomodulation (PBM) to enhance brain health, specifically glymphatic drainage and thus neurotoxic waste clearance, may make it a promising therapeutic tool against neurodegenerative diseases such as Alzheimer’s disease. This study investigates whether PBM can modulate cerebrospinal fluid (CSF) flow in 45 healthy young adults. We conducted forehead transcranial PBM (tPBM) and intranasal PBM (iPBM) at the nostril level, and measured CSF dynamics using blood-oxygenation level-dependent (BOLD) functional MRI (fMRI). Our data demonstrates 4 min of PBM-induced increases in CSF flow. Our data shows that (1) even a short PBM of 4 min can induce a change in CSF dynamics, in the form of an immediate increase in intracranial CSF volume and a reduction in CSF inflow; (2) skin melanin had a significant effect on the CSF response in tPBM, with lighter skin associated with higher responses; (3) both iPBM and tPBM displayed a dose-dependent effect on CSF dynamics in terms of a irradiance-wavelength interaction; (4) intranasal PBM (iPBM) can be used to produce a significant change in CSF dynamics that is equivalent to forehead transcranial PBM (tPBM) with a small fraction of the irradiance. The most likely explanation for the observed fMRI signal changes in CSF regions of interest for both tPBM and iPBM is an increased CSF outflow pressure due to PBM-induced vasodilation that transiently increases intracranial CSF volume and reduces net CSF inflow. This study establishes that PBM can modulate CSF flow in the healthy human brain in real time. This study also suggests that iPBM may be more efficient in CSF modulation due to the proximity to the olfactory system and the lack of melanin dependence. The influence of melanin on tPBM, the feasibility of iPBM and the dose dependence of both will require further investigation in healthy and patient populations.

## Introduction

The flow of the cerebrospinal fluid (CSF) is now known to play a vital role in brain health and disease. CSF production and movement are crucial for delivering essential nutrients and, importantly, for clearing metabolic waste products (Rasmussen et al., 2022). Specifically, CSF flow is intricately tied to glymphatic flow. The glymphatic system is the waste clearance center of the central nervous system and utilizes cerebrospinal fluid (CSF) as a carrier for toxic materials. Due to the role of CSF-interstitial fluid (ISF) exchange in glymphatic flow, CSF flow has been taken as a surrogate of glymphatic flow (Fultz et al., 2019; Rasmussen et al., 2018; Wang et al., 2023). Impairment of glymphatic flow has been implicated in neurodegenerative disorders such as Alzheimer’s disease, Parkinson’s disease, and other forms of dementia (Rasmussen et al., 2018). Recently, blood-oxygenation level-dependent (BOLD) functional MRI (fMRI) has become popularized for imaging CSF dynamics by leveraging the inflow effect (Attarpour et al., 2021; Fultz et al., 2019; Williams et al., 2023; H.-C. S. Yang et al., 2022). Moreover, fMRI-based dynamic CSF measurements have revealed that sleep, respiration and functional tasks can all modulate CSF inflow and outflow (Fultz et al., 2019; Williams et al., 2023; Yang et al., 2024; Zhong et al., 2024). However, despite its implications for improving brain health, the modulation of CSF flow by transient neuromodulation techniques has yet to be demonstrated.

In recent years, photobiomodulation (PBM) has emerged to be a promising tool for enhancing brain health. PBM refers to the use of low-intensity near-infrared light to stimulate biological processes (Mester et al., 1968). The majority of existing PBM studies have used continuous irradiation (Nairuz et al., 2024; Ramakrishnan et al., 2024; Yan et al., 2025; Yoo, 2021; Zein et al., 2018), with pulsed PBM being a more novel approach that is potentially more efficient than continuous PBM (Zein et al., 2018). PBM has been linked to increased production of adenosine triphosphate (ATP) through the oxidation of the chromophore cytochrome c oxidase (CCO) in the cellular mitochondria (Hamblin and Demidova, 2006; Pastore et al., 2000), as well as to an increased release of the potent dilator nitric oxide (NO) (Hamblin and Demidova, 2006; Poyton and Ball, 2011). There is a growing interest in harnessing PBM for glymphatic regulation. PBM-induced production of NO has been suggested as a potential mechanism for the enhanced glymphatic drainage associated with PBM therapy (Salehpour et al., 2022; Saucedo et al., 2021). In addition, previous studies have demonstrated improved amyloid clearance in Alzheimer’s mouse models, which were attributed to improved glymphatic pumping due to the effect of PBM on the astrocyte endfeet (Zinchenko et al., 2019). However, until recently, there has been no experimental confirmation in humans.

Recently, using two-channel broadband near-infrared spectroscopy (NIRS) before and after continuous tPBM, recent work by Saeed et al. was the first to demonstrate a transient tPBM-related increase in the steepness of the negative slope between hemoglobin [HbT] and free-water concentration [H_2_O_free_] in the human brain, which indirectly suggests an increased vascular-CSF coupling that increases CSF outflow (Saeed et al., 2025). However, it is unclear (i) whether PBM directly results in an increased CSF inflow or outflow that can be observed during stimulation; (ii) whether pulsed PBM could also produce an observable CSF response; (iii) if the observed effects vary with PBM dosage parameters. Specifically, PBM dosimetry is a complex topic (Cassano et al., 2019; Van Lankveld et al., 2024; Zein et al., 2018), and delivering sufficient light to the brain remains a challenge (Nairuz et al., 2024), and requires optimizing PBM stimulation parameters. PBM wavelength, frequency, and irradiance all play potentially crucial roles. Moreover, skin tone also influences light penetration, as shown in our recent work based on simulations (Van Lankveld et al., 2024) and experiments (Van Lankveld et al., 2025), but its effect on CSF modulation by tPBM is unknown. Furthermore, the effectiveness of different PBM positions, notably intranasal PBM as compared to forehead tPBM, is also unknown.

In this work, we use BOLD fMRI derived from pseudo-continuous arterial-spin labeling (pCASL) to estimate CSF dynamics. Our primary research objective was to determine whether pulsed PBM can modulate CSF dynamics in-vivo in humans. Secondarily, we want to demonstrate the influence of dose factors and skin tone on the PBM-induced CSF association. We will also compare the outcome of PBM delivered transcranially (tPBM) on the forehead and intranasally at the nostril (iPBM).

## Theory and hypothesis

### CSF dynamics and PBM

In the recent study by Saeed et al. (Saeed et al., 2025), the ratio of HbT and H_2_O_free_ in healthy adults was measured using two-band NIRS. This ratio was used as a surrogate to vascular-CSF coupling. Saeed et al. found that after 8 minutes of continuous-wave forehead tPBM, the slope characterizing H_2_O_free_ relative to HbT became more significantly negative in the prefrontal brain region. Moreover, the ability of tPBM to enhance microcirculation and vasodilation through NO release has been experimentally demonstrated (Wang et al., 2017). The vasodilation is in turn expected to translate into increased CSF outflow (Zimmermann et al., 2025). Taken together, these findings suggest that tPBM boosts the driving mechanism of CSF outflow, thereby promoting glymphatic circulation and enhancing waste clearance from the brain.

### CSF dynamics in fMRI

Translation of the NIRS findings by Saeed et al. into fMRI measurements provides us with the following hypotheses, which are further illustrated in **Figure 2**: (i) PBM is associated with an increase in the fMRI signal in CSF-related regions (except for the inflow slice), reflecting an increase in CSF volume; (ii) PBM is associated with a decrease in the fMRI signal in the inflow slice, reflecting a decrease in CSF inflow that is potentially driven by the increased intracranial CSF volume.

**Figure 2.**
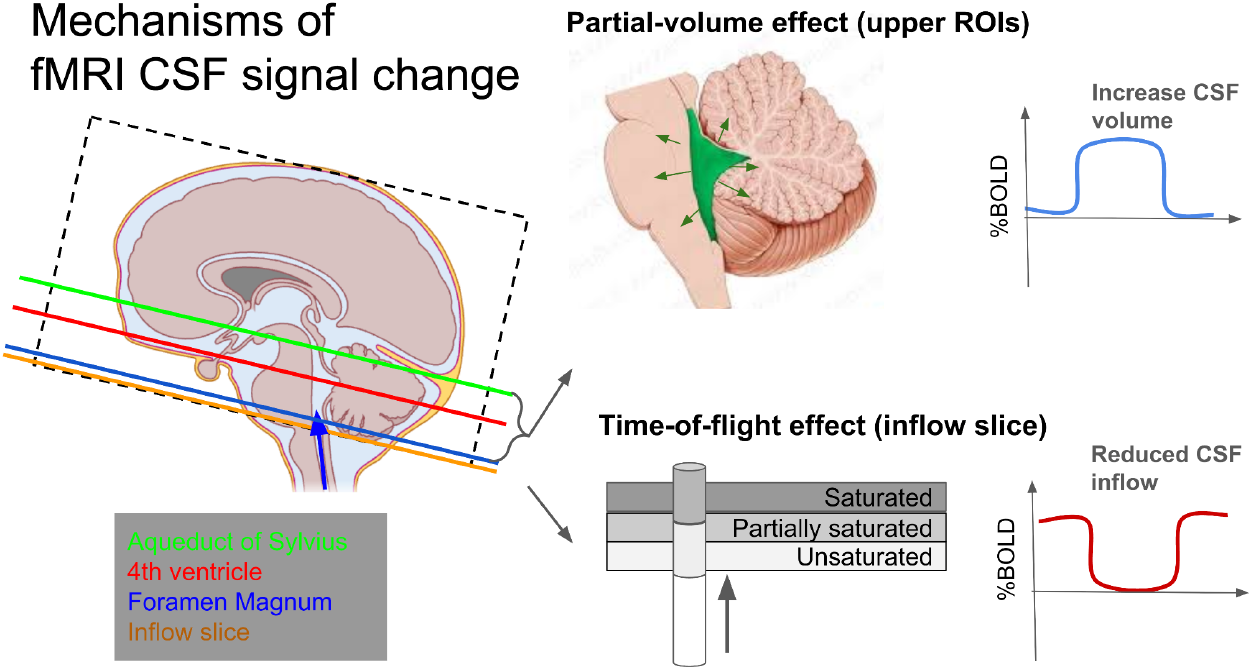
Theoretical mechanisms relating CSF volume and inflow to the fMRI signal. Translating prior findings to the fMRI space, an increase in CSF volume translates to an increase in BOLD signal in different locations along the CSF-flow pathway, namely the aqueduct, the fourth ventricle and the foramen magnum. The blue arrow indicates the direction of CSF inflow.

The basis of a CSF-volume based fMRI signal change is the partial-volume effect. The signal intensity (S) of a given fMRI voxel consists of a contribution from the CSF volume (V) and the tissue volume (1-*V*),

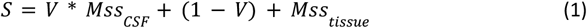

where *Mss*_CSF_ and *Mss*_tissue_ are the steady-state MRI signals associated with CSF and brain tissue, respectively. The steady-state MRI signal intensity (*Mss*) for each compartment (CSF or tissue) is defined by the Ernst equation,

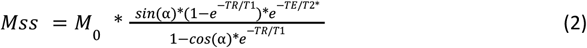

where *M*_*0*_ is the equilibrium magnetization, α is the flip angle, T1 is the longitudinal relaxation time, T2* is the transverse relaxation time, TR is the repetition time and TE is the echo time. The fMRI signal change (dS) associated with CSF volume change (dV) depends heavily on the composition of the voxel at baseline, as shown in **Fig. 3**. In this case, we simulated grey-matter tissue, with the following parameters for 3 Tesla: T1 = 1.3 s (Wansapura et al., 1999), T2* = 40 ms (Barry and Smith, 2019), TR = 4.5 s (representative of our pCASL MRI acquisition), TE = 30 ms. Moreover, *V* becomes *V*\*(1+*dV*), and it is assumed that the tissue fraction becomes 1-*V*\*(1+*dV*).

**Figure 3.**
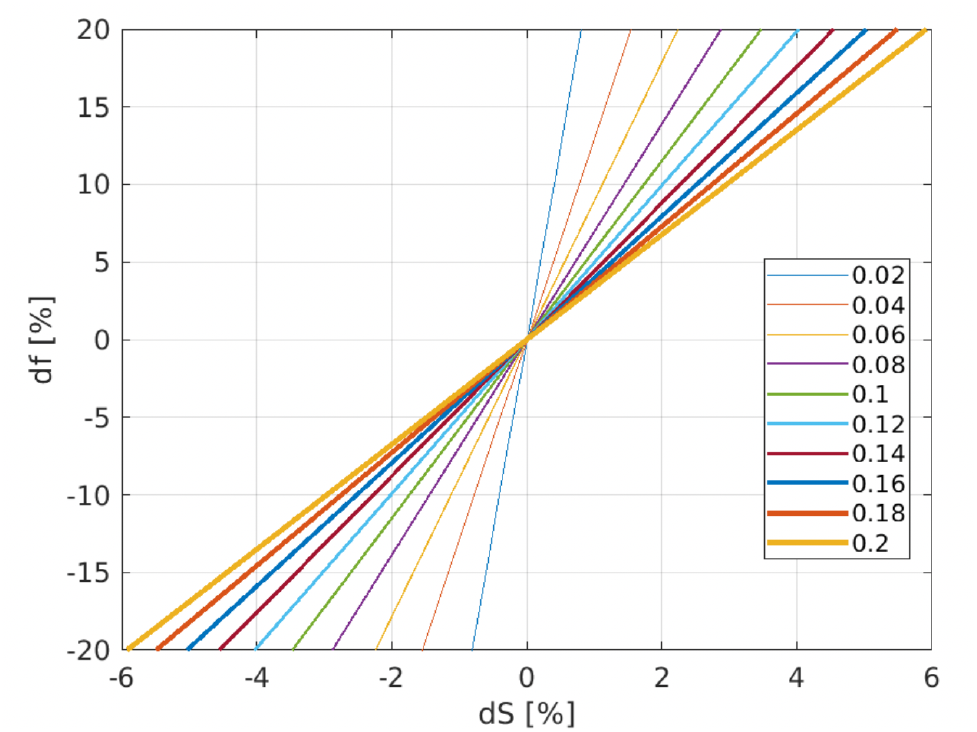
Theoretical prediction of fMRI signal change (dS) due to a change in voxelwise CSF volume fraction (df). The plots are shown for different levels of baseline CSF fraction (indicated in the legend, from 2% to 20%).

Furthermore, using the well-defined inflow equations, we can estimate the approximate velocity change that corresponds to this level of CSF-related fMRI signal change (Gao and Liu, 2012),

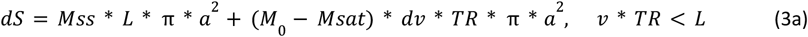

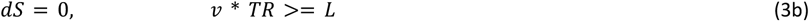

where v is the flow velocity, L is the slice thickness and a is the radius of the CSF ROI in plane. The equation is visualized in **Fig. 4** for different values of *a*. At an inflow decrease of >100%, CSF inflow reverses to CSF outflow completely.

**Figure 4.**
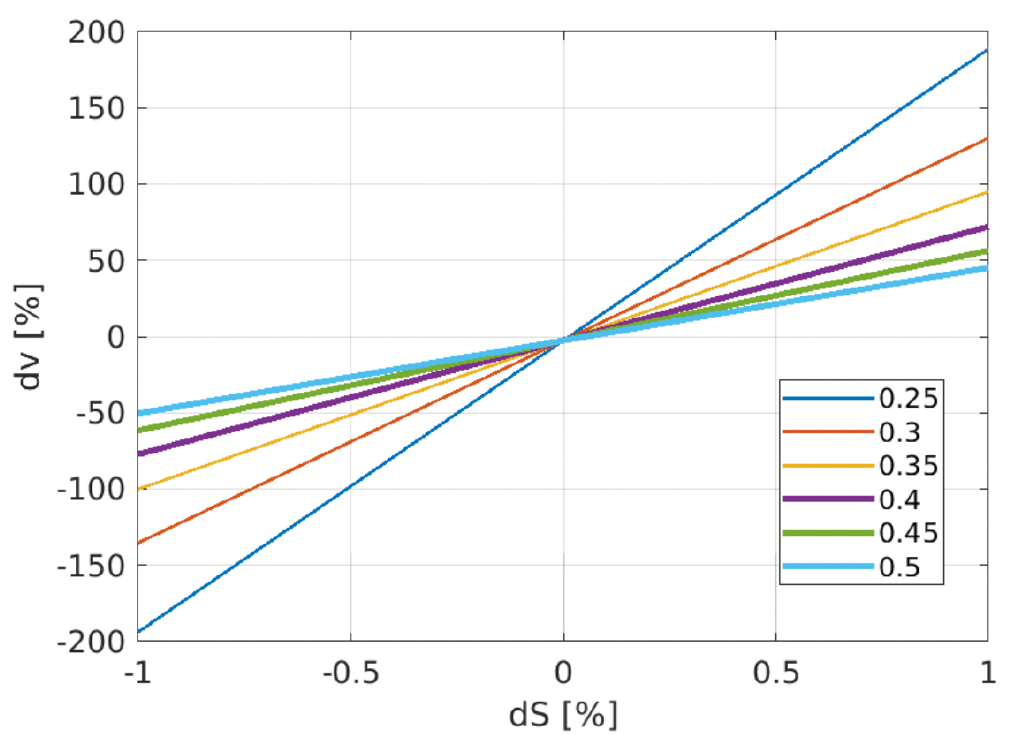
Theoretical prediction of fMRI signal change (dS) due to a change in CSF inflow velocity (dv). The lines represent the theoretical conversion between dS and dv in the inflow ROI, assuming an inflow-regional radius (a) ranging from 0.25 to 0.5 mm (labeled in the legend), with a slice thickness of 4 mm and a TR of 4.5 s. A negative value for dv represents a reduction in inflow velocity.

**Figure 5.**
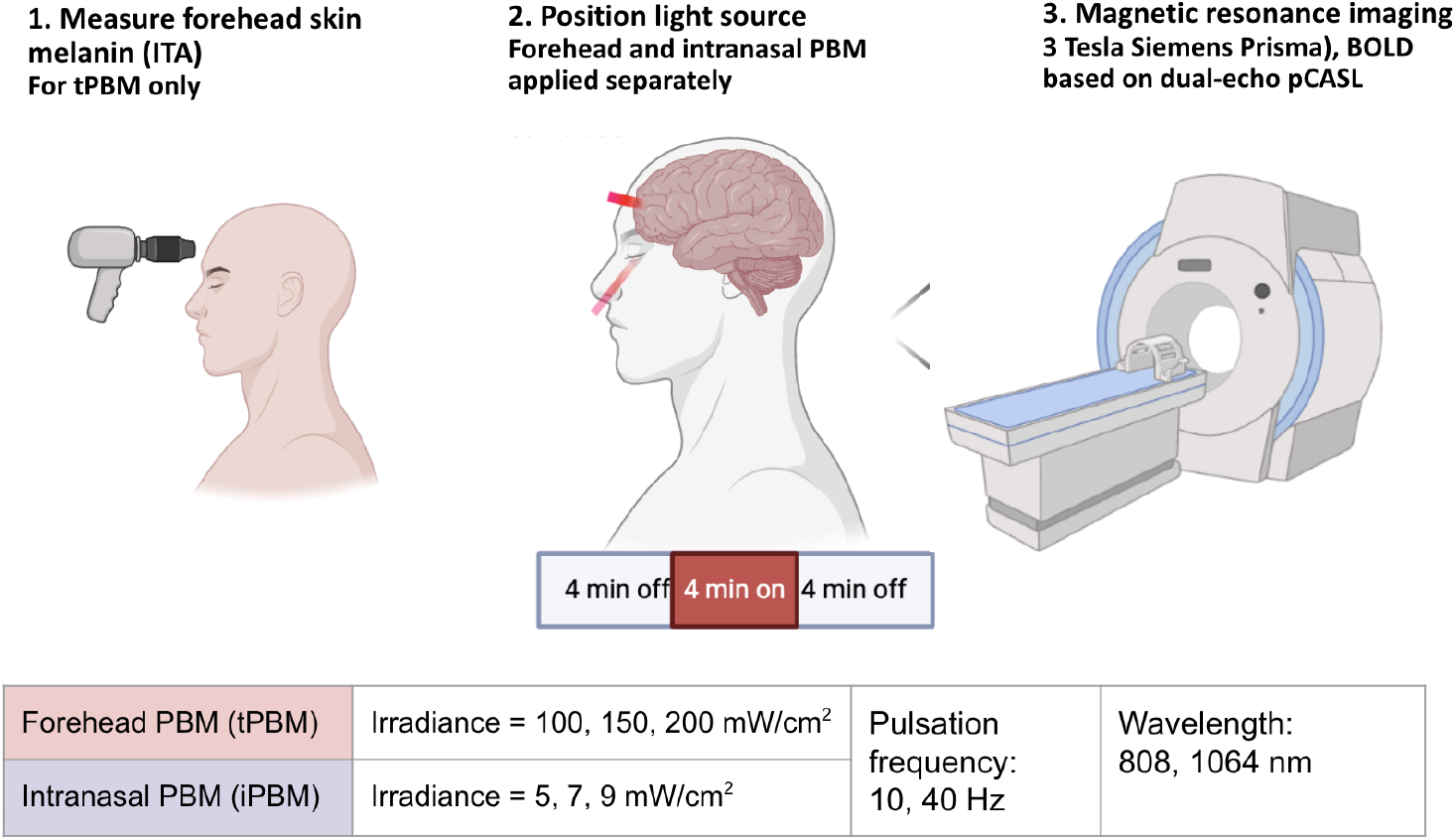
Summary of experimental set-up.

### PBM dose dependence

In theory, light deposition inside the brain depends on light wavelength, irradiance, positioning (Cassano et al., 2019; Zein et al., 2018), and in the case of pulsed PBM, the pulsation frequency (Kim et al., 2017). Each parameter can be adjusted to impact the overall energy accumulation in tissue. Previous research, based on cultured cortical neurons, suggests that the peak PBM response happens when the energy induced by the light source (i.e. the irradiance) reaches 0.3-3 J/cm^2^ in the specified brain region (Huang and Hamblin, 2019). Past simulation studies using the Monte Carlo method by ourselves and others demonstrated that the 808 nm wavelength is likely to produce significantly higher fractional energy deposition than longer or shorter wavelengths in tPBM (Cassano et al., 2019; Li et al., 2017; Van Lankveld et al., 2024). Moreover, power deposition should scale linearly with irradiance (Van Lankveld et al., 2024). Notably, our simulations showed that lighter skin tones allowed higher energy accumulation. Moreover, the 810 nm wavelength for tPBM and 1064 nm wavelength for iPBM produced the highest cortical energy accumulation, which was linearly correlated with optical power density, but these variations could be overridden by a difference in skin tone in the tPBM case.

Based on these simulation-based findings, we hypothesize that (i) a wavelength of 808 would be associated with the highest CSF response; (ii) the response should scale positively with irradiance.

## Methods

### Participants

We studied 45 healthy young adults (aged 20-32, 23 M / 22 F). Participants were screened prior to their participation to ensure there was no history of neurological or physiological disorders, malignant disease, or the use of medications that could have influenced the study. All experiments were conducted in accordance with REB guidelines, and all participants signed informed consent, approved by the Baycrest Research Ethics Board (REB).

### Melanin measurement

We targeted an equal representation of three skin colour categories (light, intermediate and dark). The individual topography angle (ITA) (related to skin colour) of each participant was measured using a CM-600D Spectrophotometer (Konica Minolta, Tokyo, Japan). The ITA is calculated as (Eq. 4),

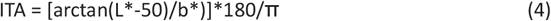

where *L* represents luminance ranging from black (0) to white (100), and *b*\* ranging from yellow to blue (12). The higher the ITA, the lighter the skin. The ITA skin colour types are typically classified into six ranges: very light (>55°), light (41° to <55°), intermediate (28° to <41°), tan (10° to <28°), brown (−30° to <10°) and dark (<-30°). Zero and White Calibration were performed before each use to ensure accuracy of the calculations. Six 8 mm diameter measurement samples were taken of the right forehead of each subject prior to tPBM laser placement and the average was computed into the individual melanin groupings.

### PBM protocols

Based on existing PBM literature the stimulation parameters chosen for this study consisted of two wavelengths (808 nm and 1064 nm), two pulsation frequencies (10 Hz and 40 Hz), and three optical power densities for each positioning (**Table 1**). The light sources are MR-compatible Class-3 lasers (courtesy: Vielight Inc., Toronto, Canada) each with a fixed wavelength and connected to a waveform generator to produce the desired duty cycle. The light is delivered via the forehead (tPBM) and at the nostril (iPBM).

**Table 1.**
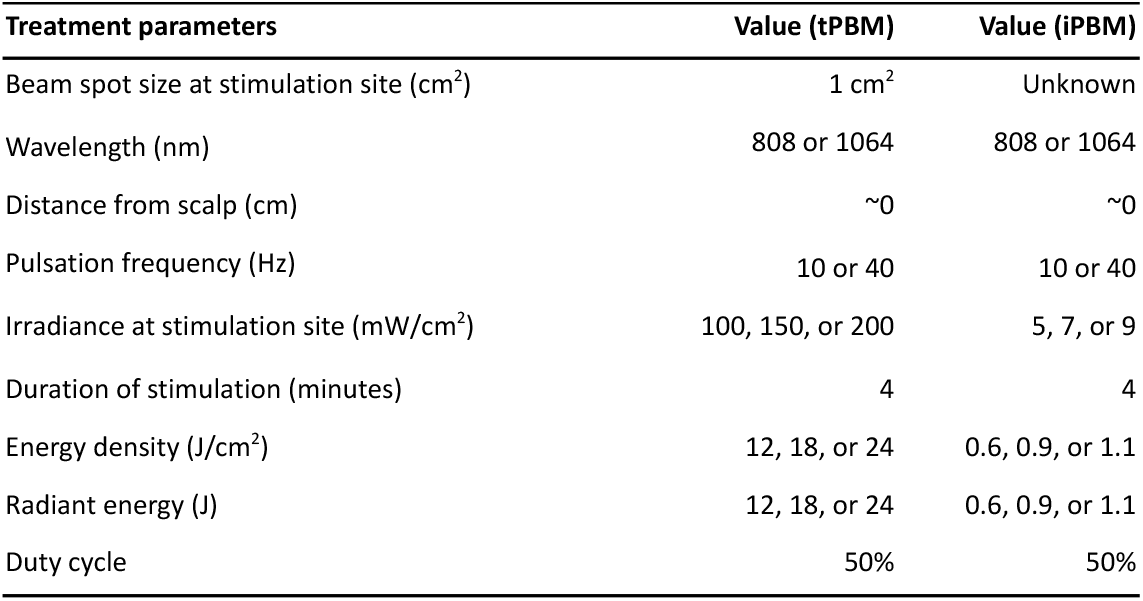
Treatment parameters.

| Treatment parameters | Value (tPBM) | Value (iPBM) |
| --- | --- | --- |
| Beam spot size at stimulation site (cm<sup>2</sup>) | 1 cm<sup>2</sup> | Unknown |
| Wavelength (nm) | 808 or 1064 | 808 or 1064 |
| Distance from scalp (cm) | ~0 | ~0 |
| Pulsation frequency (Hz) | 10 or 40 | 10 or 40 |
| Irradiance at stimulation site (mW/cm<sup>2</sup>) | 100, 150, or 200 | 5, 7, or 9 |
| Duration of stimulation (minutes) | 4 | 4 |
| Energy density (J/cm<sup>2</sup>) | 12, 18, or 24 | 0.6, 0.9, or 1.1 |
| Radiant energy (J) | 12, 18, or 24 | 0.6, 0.9, or 1.1 |
| Duty cycle | 50% | 50% |

Each fMRI scan was 12 minutes long and followed a block design (OFF-ON-OFF) stimulation paradigm (see **Fig. 6**): 4 minutes of baseline, 4 minutes of PBM stimulation (DURING), and 4 minutes of post-stimulation (POST). This duration was chosen as it is within the range used in PBM literature and allows us to monitor both online and offline effects (Dmochowski et al., 2020; Wang et al., 2019). Combining the 4-minute baseline period, the 4-minute post-stimulation period, and the rest between consecutive scans, there was a minimum of 20 minutes between periods of active PBM stimulation.

**Figure 6.**
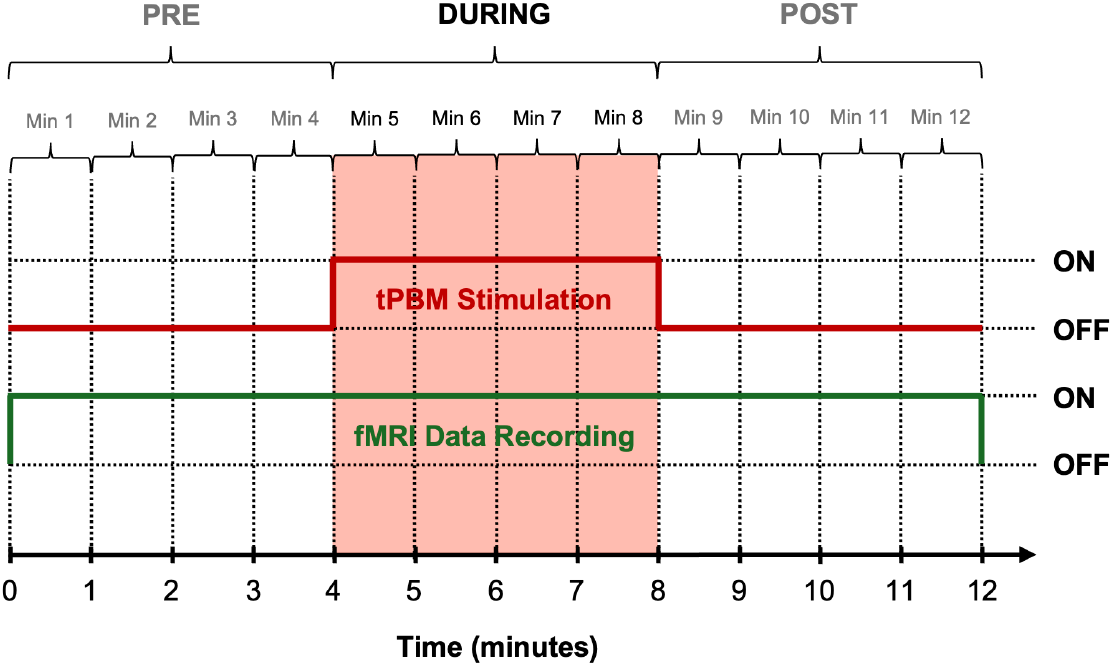
PBM timeline diagram. The green line represents the fMRI data recording period, and the pink shaded block shows the PBM-ON period. The same timing is used for iPBM and tPBM.

To thoroughly evaluate the effects of these different PBM light parameters within a reasonable experimental duration, each participant was randomly assigned to one of three protocols which determined the exact parameter combinations they were exposed to during stimulation for all four EEG scans (**Table 2**). Prior to the experiments, all irradiance levels were tested on a small group of healthy subjects to ensure the absence of all thermal sensation. Moreover, each subject was asked about heating sensations but none was reported. As we ensured that PBM was not associated with any heating or sound, the participants were ignorant of when the light was on or off. A breakdown of the subject distributions across protocols is also shown in **Table 2**. Furthermore, a breakdown of subject numbers that also incorporates melanin categories is shown in **Table 3**. A breakdown of the ITA distribution of samples across melanin categories is shown in **Figure 7**.

**Table 2.**
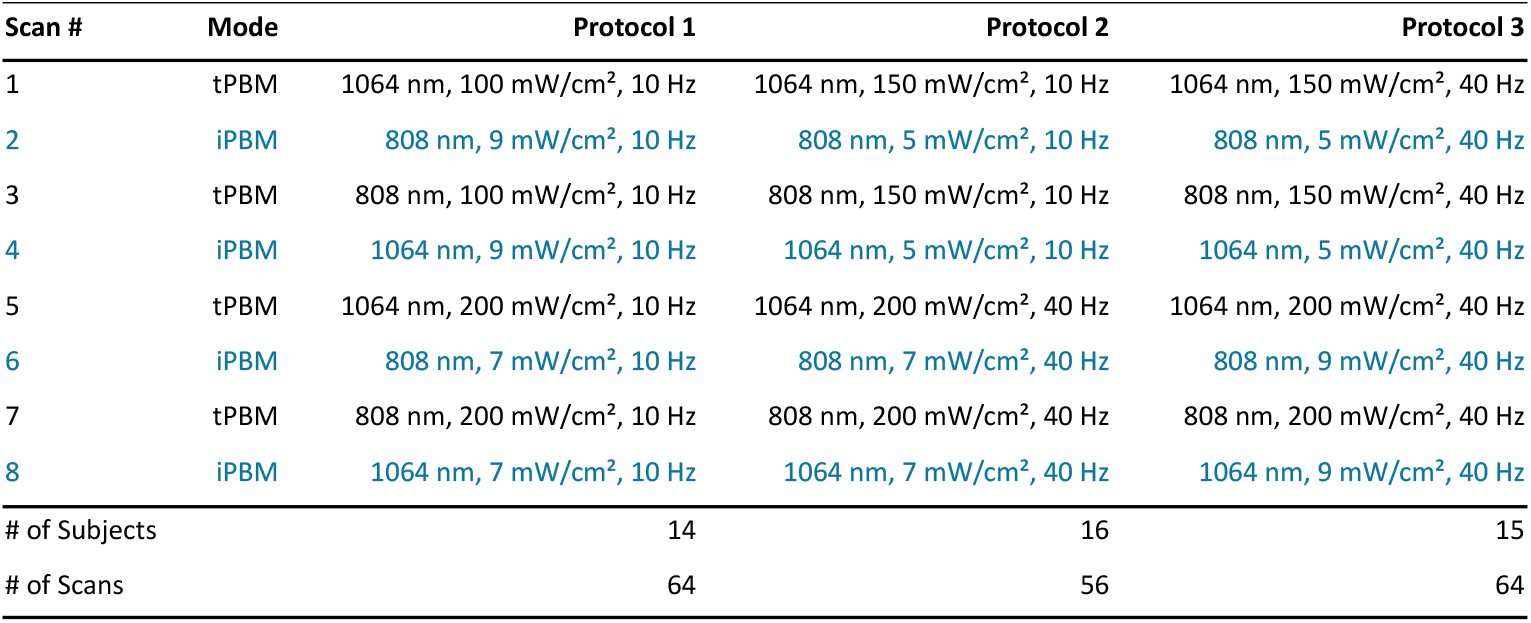
12-min tPBM and iPBM scan distribution by protocol.

| Scan # | Mode | Protocol 1 | Protocol 2 | Protocol 3 |
| --- | --- | --- | --- | --- |
| 1 | tPBM | 1064 nm, 100 mW/cm<sup>2</sup>, 10 Hz | 1064 nm, 150 mW/cm<sup>2</sup>, 10 Hz | 1064 nm, 150 mW/cm<sup>2</sup>, 40 Hz |
| 2 | iPBM | 808 nm, 9 mW/cm<sup>2</sup>, 10 Hz | 808 nm, 5 mW/cm<sup>2</sup>, 10 Hz | 808 nm, 5 mW/cm<sup>2</sup>, 40 Hz |
| 3 | tPBM | 808 nm, 100 mW/cm<sup>2</sup>, 10 Hz | 808 nm, 150 mW/cm<sup>2</sup>, 10 Hz | 808 nm, 150 mW/cm<sup>2</sup>, 40 Hz |
| 4 | iPBM | 1064 nm, 9 mW/cm<sup>2</sup>, 10 Hz | 1064 nm, 5 mW/cm<sup>2</sup>, 10 Hz | 1064 nm, 5 mW/cm<sup>2</sup>, 40 Hz |
| 5 | tPBM | 1064 nm, 200 mW/cm<sup>2</sup>, 10 Hz | 1064 nm, 200 mW/cm<sup>2</sup>, 40 Hz | 1064 nm, 200 mW/cm<sup>2</sup>, 40 Hz |
| 6 | iPBM | 808 nm, 7 mW/cm<sup>2</sup>, 10 Hz | 808 nm, 7 mW/cm<sup>2</sup>, 40 Hz | 808 nm, 9 mW/cm<sup>2</sup>, 40 Hz |
| 7 | tPBM | 808 nm, 200 mW/cm<sup>2</sup>, 10 Hz | 808 nm, 200 mW/cm<sup>2</sup>, 40 Hz | 808 nm, 200 mW/cm<sup>2</sup>, 40 Hz |
| 8 | iPBM | 1064 nm, 7 mW/cm<sup>2</sup>, 10 Hz | 1064 nm, 7 mW/cm<sup>2</sup>, 40 Hz | 1064 nm, 9 mW/cm<sup>2</sup>, 40 Hz |
| # of Subjects |  | 14 | 16 | 15 |
| # of Scans |  | 64 | 56 | 64 |

**Table 3.** Participant distribution by protocol and melanin group.

|  | Protocol 1 | Protocol 2 | Protocol 3 |
| --- | --- | --- | --- |
| **Melanin Group 1** | 5 | 5 | 4 |
| **Melanin Group 2** | 6 | 5 | 6 |
| **Melanin Group 3** | 5 | 4 | 6 |
| # of Subjects | 14 | 16 | 15 |
| # of Scans | 64 | 56 | 64 |

**Figure 7.**
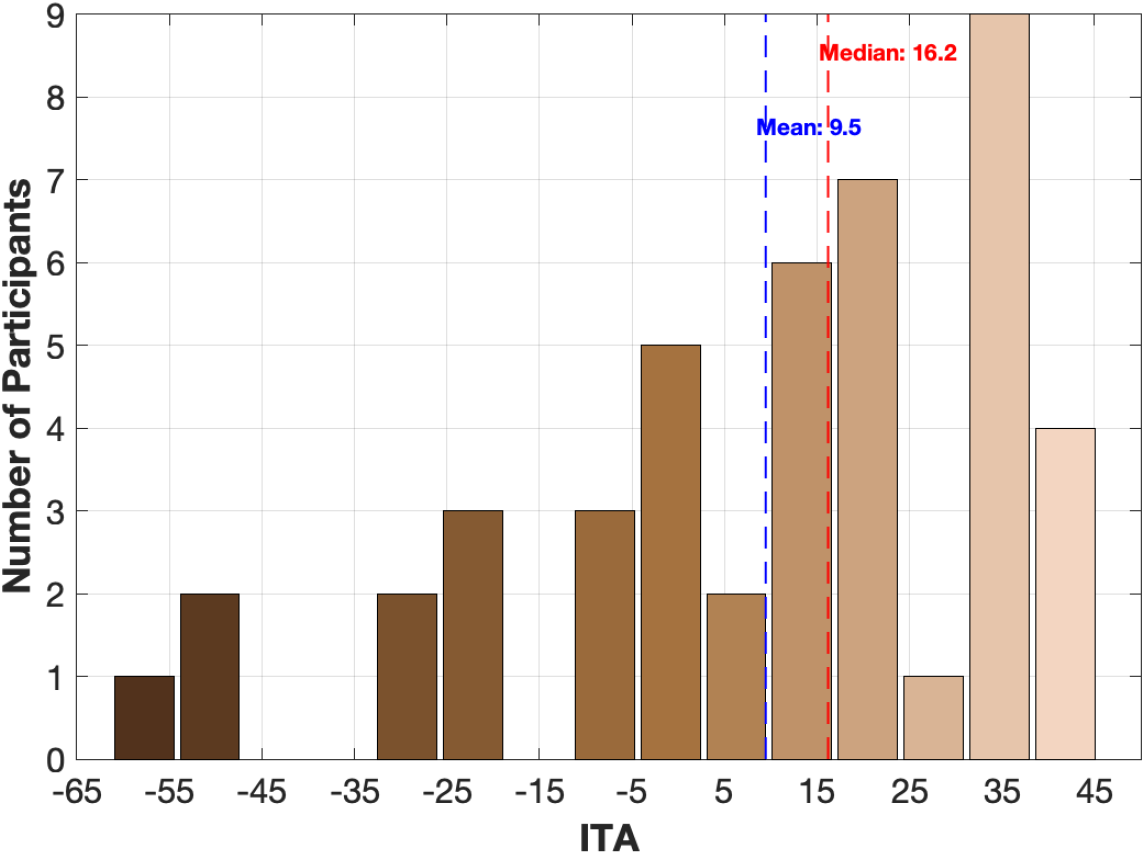
Distribution of participants across skin tones. The colours represent the skin tone associated with different individual topography angles (ITA) for the different participants. The mean and median ITA values are indicated in blue and red, respectively.

**Figure 8.**
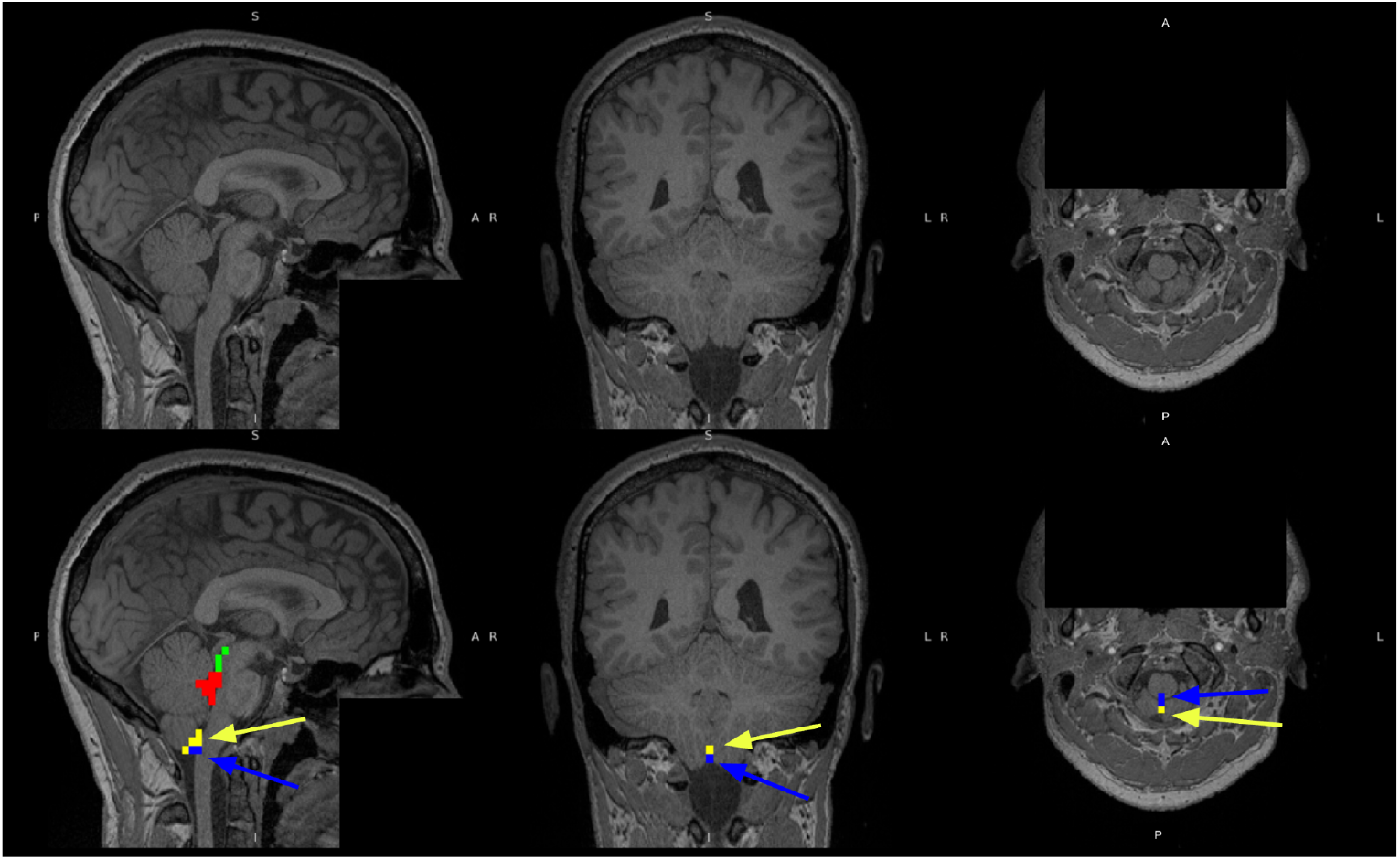
Regions-of-interest (ROI) definitions. The defaced high-resolution T1 anatomical images in a sample subject are shown in the top row for reference. The regions of interest defined in fMRI space are shown in the bottom flow for the same subject. Yellow corresponds to the inflow-slices (3 slices labelled, only the lowest slice for each rs-fMRI session was included in analysis), red the 4th ventricle, green the aqueduct, and blue the foramen magnum.

### MRI acquisition

The images were acquired using a Siemens Prisma 3 Tesla System (Siemens, Erlangen, Germany), which employed 20-channel phased-array head coil reception and body coil transmission. We acquired T1-weighted 3D anatomical images (sagittal, 234 slices, 0.7 mm isotropic resolution, TE = 2.9 ms, TR = 2240 ms, TI = 1130 ms, flip angle = 10°) for positioning and CSF-region segmentation. We acquired BOLD fMRI data with dual-echo (DE) pseudo-continuous arterial spin labelling (pCASL) (courtesy of Danny J. J. Wang, University of Southern California) for recording CSF-related BOLD dynamics (TR = 4.5 s, TE1 = 9.8 ms, TE1 = 30 ms, post-labelling delay = 1.5 s, labelling duration = 1.5 s, flip angle (*α*) = 90°, 3.5 mm isotropic resolution, 35 slices, slices gap = 25%, scanning time = 4 minutes), in which BOLD dynamics were extracted from the 2^nd^ TE data by surround averaging after low-pass filtering (Halani et al., 2015; Tak et al., 2014).

### Data analysis

#### fMRI data preprocessing

Preprocessing was performed using a custom script that employed the FMRIB Software Library (FSL) for brain extraction, motion correction, distortion correction, slice timing, temporal and spatial outlier removal, drift correction and rigorous scalp stripping. A 0.1Hz low pass filter was also applied to the CBF data as well as thresholding. The first five volumes of both BOLD and CBF were rejected to allow the brain to enter a steady state.

#### Regions-of-interest definition

Using the T1 anatomical image for each participant, we manually segmented the fourth ventricle, aqueduct, and foramen magnum regions of interest (ROIs). Additionally, to maximize the in-flow effect, we manually identified a CSF ROI in the lowest slice for each fMRI scan.

#### General-linear model analysis

The mean time courses for all ROIs were submitted to general linear model (GLM) analyses to assess the presence of significant PBM response. Each mean time course was regressed against the timing waveform (**Fig. 6**), and all regression coefficients were thresholded at the p<0.05 level. These were then submitted to false-discovery rate (FDR) correction using the Benjamini-Hochberg method (Benjamini and Hochberg, 1995).

#### Linear mixed-effects analysis

All significant regression coefficients were submitted to a three-stage Linear Mixed Effects (LME) modeling pipeline. Variables considered for fixed effects included wavelength, irradiance, pulsation frequency and ITA as well as their interactions. With the exception of the ITA, all fixed-effect variables are considered categorical. The subject ID was used as the random-effect variable.

In the first stage, we used stepwiselm in Matlab (Natick, MA, USA) to remove irrelevant variables from consideration. That is, starting from a constant, variables (and their interaction terms) were gradually added to the model in a stepwise manner and the significance of the effect size were determined. If the effect size of a given variable failed to meet the p < 0.05 criterion, then that variable was removed. In this way, a preliminary model was determined for further testing. Melanin was not relevant for iPBM, so ITA was not included as a dependent variable in the LMEs for iPBM.

In the second stage, a final LME model was constructed using all variables that survived the stepwise regression in the first stage, and the surviving effects are considered significant if associated FDR-corrected p-values were lower than 0.05.

## Results

### CSF response in tPBM

#### Response for different regions of interest

Group-mean fMRI responses for tPBM in the CSF ROIs are shown in **Figure 9**. In each column, the plots are averaged across all values for the other two variables. These time courses serve to prove the feasibility of observing the real-time CSF signal response to PBM.

**Figure 9.**
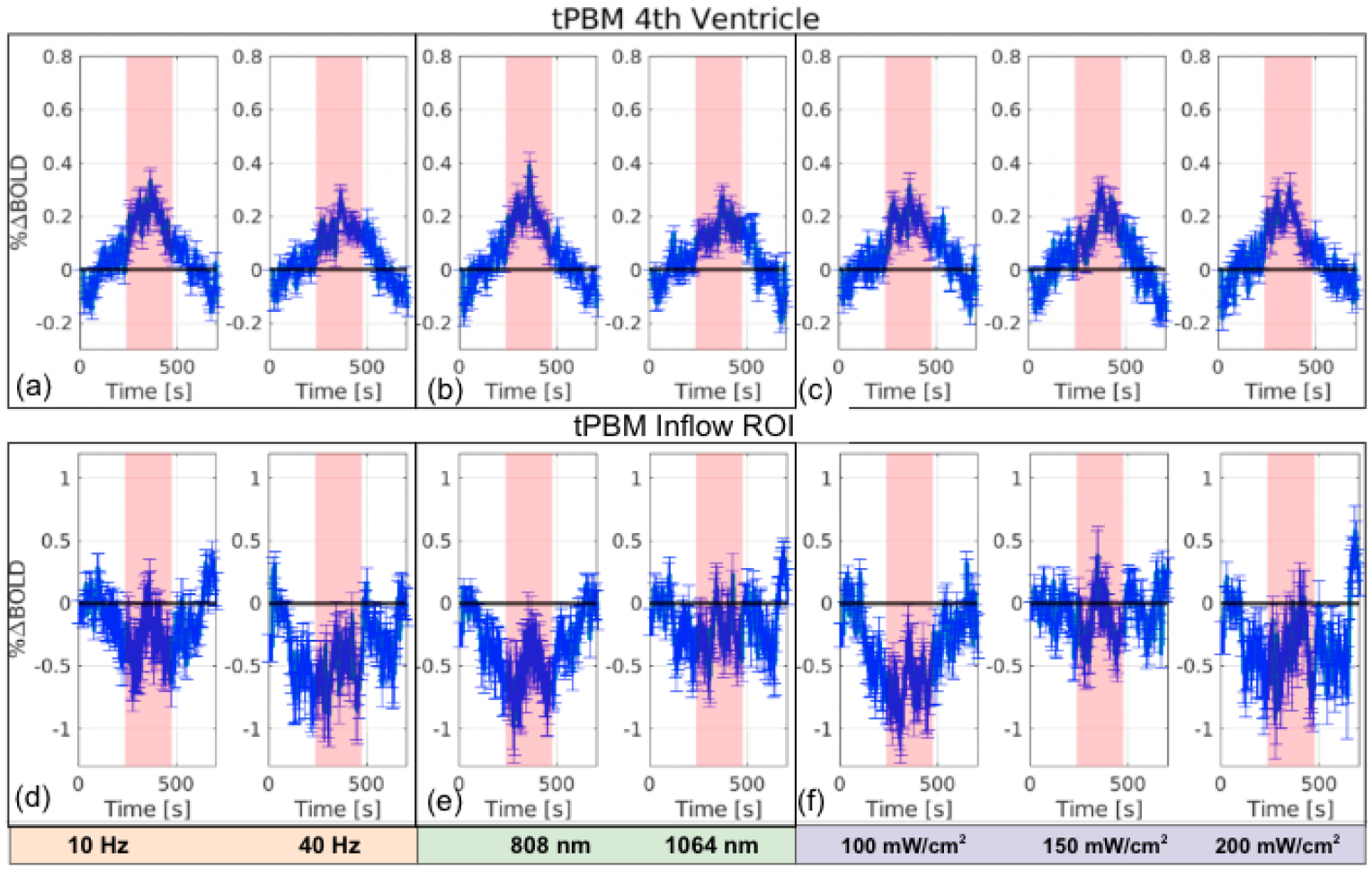
tPBM CSF responses in the fourth ventricle and the inflow ROI. The Mean BOLD fMRI time courses are shown in blue. In each average, the time courses from all scans of all participants are included, and grouped by the variable of interest only (collapsing across all other variable dimensions). (a) and (d) are grouped by frequency, (b) and (e) grouped by wavelength and (c) and (f) grouped by irradiance. The error bars represent the standard error.

The 4th ventricle displayed a signal increase during the PBM-on period that quickly dissipated as the laser was turned off, consistent with a transient increase in CSF volume. Similar responses are seen in other upper-CSF ROIs, shown in **Fig. S1** in Supplementary Materials. The change in fMRI signal range averages at ∼0.3%, which, according to theoretical predictions in **Fig. 3**, can correspond to a CSF volume increase of between 3% and > 20%. The Inflow ROI displayed the opposite trend for most of the settings, consistent with a reduction in CSF inflow during tPBM. The change in fMRI signal range averages at ∼-0.6%, which, according to theoretical predictions in **Fig. 4**, can correspond to a CSF inflow decrease of between 30 and 120%.

#### Skin-tone dependence

The effect of melanin is summarized in **Table 4**. As the inflow slice showed a negative CSF response, the absolute value was taken when performing the LME. It was found that the ITA has a significant positive effect on the CSF signal in the aqueduct and the inflow slices. That is, fairer skin (higher ITA) is associated with greater CSF response to tPBM. This effect was not apparent in the Foramen of Monro and the 4th ventricle.

**Table 4.** Effect sizes and p values (subject as random effect).

| tPBM | Irradiance |  | Wavelength |  |  | Frequency |  |  | ITA |
| --- | --- | --- | --- | --- | --- | --- | --- | --- | --- |
|  | 808 nm | 1064 nm | 100 mW/cm<sup>2</sup> | 150 mW/cm<sup>2</sup> | 200 mW/cm<sup>2</sup> | 100 mW/cm<sup>2</sup> | 150 mW/cm<sup>2</sup> | 200 mW/cm<sup>2</sup> |  |
| **Aqueduct** | 100<150 mW/cm<sup>2</sup><br>(*t*(162) = 3.20,<br>*p* = 0.0017, *CI* = [0.37, 1.57]) |  | – | – | – | – | 10Hz > 40Hz<br>(*t*(162) = 2.25,<br>*p* = 0.030, *CI* = [0.10, 1.52]) | – | +ve<br>(*t*(162) = 2.50,<br>*p* = 0.0013, *CI* = [0.0014, 0.012]) |
| **Foramen** | – | – | – | – | – | – |  |  | – |
| **4th Ventricle** | – | – | – | – | – | – |  |  | – |
| **Inflow** | 150>(100, 200) mW/cm<sup>2</sup><br>(*t*(162) = 2.83,<br>*p* = 0.0052, *CI* = [0.26, 1.48]) | 150<(100, 200) mW/cm<sup>2</sup><br>(*t*(162) = 2.56,<br>*p* = 0.011, *CI* = [0.18, 1.25]) | – | – | – | – |  |  | +ve<br>(*t*(162) = 1.82,<br>*p* = 0.01, *CI* = [-0.0010, 0.013]) |

#### Dose dependence

The dose dependence in the LME was revealed as interactions between irradiance and wavelength, and between irradiance and pulsation frequency (**Table 4**). Specifically, in the inflow ROI, the CSF response was maximized at 150 mW/cm^2^ but only at 808 nm; at 1064 nm, 150 mW/cm^2^ was associated with the minimal CSF response. Moreover, 10 Hz produced a larger CSF response in the cerebral aqueduct than 40 Hz, but only observable at 150 mW/cm^2^. There was no interaction between wavelength and pulsation frequency. These effects were also not apparent in the foramen magnum and the fourth ventricle.

### CSF response in iPBM

#### Response for different regions-of-interest

The fMRI-based CSF responses for iPBM are very similar to those observed for tPBM (**Fig. 10)**. The 4th ventricle displayed a signal increase during the PBM-on period, with similar responses seen in other upper-CSF ROIs, shown in **Fig. S2** in Supplementary Materials. The Inflow ROI displayed the opposite trend for most of the settings, consistent with a reduction in CSF inflow during iPBM.

**Figure 10.**
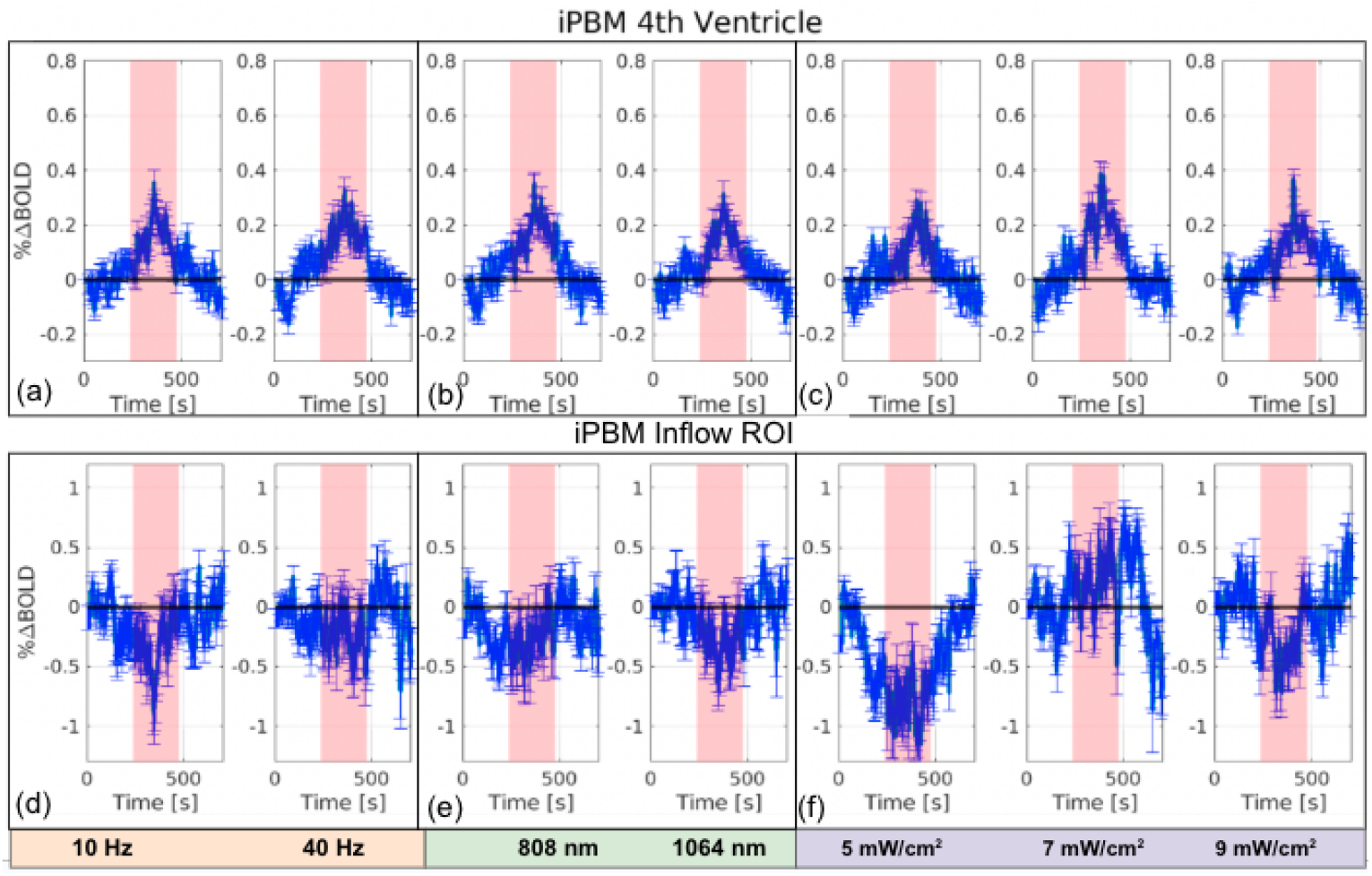
iPBM CSF responses in the fourth ventricle and the inflow ROI. The Mean BOLD fMRI time courses are shown in blue. In each average, the time courses from all scans of all participants are included, and grouped by the variable of interest only (collapsing across all other variable dimensions). (a) and (d) are grouped by frequency, (b) and (e) grouped by wavelength and (c) and (f) grouped by irradiance. The error bars represent the standard error.

#### Dose dependence

As seen in **Fig. 10**, all irradiance levels except 7 mW/cm^2^ elicited a decrease in inflow BOLD. Moreover, as shown in **Table 5**, we found a wavelength-fluence interaction that follows a biphasic irradiance dependence. That is, at 808 nm, 7 mW/cm^2^ produced the least CSF response, whereas at 1064 nm, 7 mW/cm^2^ produced the largest CSF response. No significant dependence on pulsation frequency and no other significant interactions were found.

**Table 5.** Effect sizes and p values (subject as random effect).

| iPBM | Irradiance |  | Wavelength | Frequency |
| --- | --- | --- | --- | --- |
|  | 808 nm | 1064 nm |  |  |
| Aqueduct | — |  | — | — |
| Foramen | — |  | — | — |
| 4th Ventricle | — |  | — | — |
| Inflow | $7 < (5, 9) \text{ mW/cm}^2$<br>$(t(162) = 1.93,$<br>$p = 0.04, CI = [0.010, 1.01])$ | $7 > (5, 9) \text{ mW/cm}^2$<br>$(t(162) = 2.22,$<br>$p = 0.02, CI = [0.086, 1.49])$ | — | — |

### tPBM vs. iPBM

In this study, the energy deposition of the iPBM protocol was on average only 3-5% that of the tPBM protocol (**Table 1**). However, as seen in **Figure 9** and **10**, the CSF response for the two protocols are very similar. The CSF response from both iPBM and tPBM showed a significant positive ITA dependence, and both showed a biphasic behaviour in terms of irradiance. However, tPBM and iPBM exhibit different dose dependencies. Specifically, as seen in **Table 4** and **Table 5**, while the intermediate irradiance of 150 mW/cm^2^ resulted in peak CSF response for tPBM using 808 nm light, the intermediate irradiance of 7 mW/cm^2^ resulted in peak CSF response for iPBM using 1064 nm light. Conversely, when the wavelength was swapped to 1064 nm, the irradiance level associated with the peak CSF response became associated with the minimum CSF response (significantly lower than that of the highest or lowest irradiance values); this was common to both iPBM and tPBM. Lastly, unlike tPBM, iPBM’s CSF response does not significantly differ by pulsation frequency.

## Discussion

In this study, we used BOLD fMRI as a surrogate of CSF volume and inflow, as has been demonstrated numerous times in previous work (Attarpour et al., 2021; Fultz et al., 2019; Williams et al., 2023; H.-C. S. Yang et al., 2022), and observed a significant real-time response in both CSF flow and volume in our group of healthy young adults. Our analysis revealed that (1) even a short stimulation with PBM can induce a change in CSF dynamics, in the form of an immediate increase in intracranial CSF volume and a reduction in CSF inflow, consistent with our hypothesis; (2) iPBM can be used to produce a significant change in CSF dynamics that is comparable with forehead tPBM with a small fraction of the irradiance, which is unexpected and contrary to our hypothesis that the brain response would be maximized at the highest irradiance value, given that energy deposition scaled with irradiance in our simulations (Van Lankveld et al., 2024); (3) both iPBM and tPBM displayed a dose-dependent effect on CSF dynamics in terms of an irradiance-wavelength interaction, which was also not predictable from our hypothesis; (4) skin melanin had a significant effect on the CSF response in tPBM, with lighter skin associated with higher responses, consistent with our hypothesis.

### Potential Mechanisms

PBM is known to increase cerebral blood flow, tissue oxygenation, and cognitive function (Hamblin, 2018). PBM’s mechanisms include stimulating mitochondrial function, reducing inflammation, and protecting against apoptosis (Hamblin, 2018; Nairuz et al., 2024). In Alzheimer’s disease models, PBM improves typical pathologies like memory loss and amyloid plaques, while regulating glial cell polarization and preserving mitochondrial dynamics (L. Yang et al., 2022). PBM is also known to enhance brain waste clearance by improving blood-brain barrier permeability and relaxing lymphatic vessels (Salehpour et al., 2022). Recent animal studies have reported that tPBM using a 1267-nm laser or 1065-nm LED can activate meningeal lymphatic vessels (MLVs) for enhanced glymphatic clearance of beta-amyloid from the mouse brain (Semyachkina-Glushkovskaya et al., 2023; Zinchenko et al., 2019). tPBM has also been shown to enhance the drainage system of the animal brain during sleep (Semyachkina-Glushkovskaya et al., 2024). Furthermore, PBM has been shown to enhance recruitment of microglia surrounding amyloid plaques in the rat model, potentially improving Aβ clearance (L. Yang et al., 2022).

In our study, we found an increase in BOLD fMRI signal in upper CSF ROIs during both tPBM and iPBM that is consistent with an increase in CSF volume. We also found a decrease in BOLD fMRI signal in the inflow ROI during both tPBM and iPBM. This decrease is consistent with a decrease in CSF inflow, as demonstrated in the Theory section. These findings are both consistent with an increased CSF drainage out of the brain as a direct result of PBM. These findings extend existing work, which is primarily based on forehead tPBM (Nairuz et al., 2024; Ramakrishnan et al., 2024; Semyachkina-Glushkovskaya et al., 2023; Yan et al., 2025; Yoo, 2021), to intranasal positioning. These findings also provide important evidence confirming a reduced CSF inflow that is likely caused by CSF outflow.

When intracranial pressure increases due to a rise in brain tissue volume or blood volume, the body’s immediate response is to lower the pressure by displacing the CSF, as the CSF is the most readily displaceable component (Mokri, 2001). As the volume of brain tissue or blood increases, CSF can be expelled through the intracranial space and into the spinal subarachnoid space, which can account for the observed decrease in CSF inflow effects. During this process, the volume of structures such as the cerebral aqueduct and fourth ventricle can transiently increase in response to the increased fluid pressure. These blood-volume changes and the pulsatile flow within the cerebral aqueduct can be visualized and quantified using MRI techniques like phase-contrast cine MRI (Battal et al., 2011; Oner et al., 2017). Moreover, transient non-cardiac blood-volume changes such as that produced by anesthesia or hypercapnia have also been observed to expel CSF (Zimmermann et al., 2025).

Importantly, our findings are consistent with recent findings using dual-band NIRS which found an increase in the ratio of Δ[H_2_O_free_] to Δ[HbT] after tPBM (Saeed et al., 2025). Concurrently, the positive slopes between changes in oxidized CCO concentration (Δ[oxiCCO]) and Δ[H_2_Ofree] were enhanced post-tPBM, consistent with an increase in CCO oxidation that is well-known to result from PBM. The increased [oxiCCO] is also consistent with previously reported vasodilation (Shahdadian et al., 2024; Wang et al., 2017). Taken together, the aforementioned increase in ATP production, cerebral blood flow/volume and increase in CSF outflow are potentially coupled. Thus, the proposed mechanism of PBM-induced CSF flow alteration can be summarized as a *NIR-induced vasodilation causing an increased CBV that promotes increased aggregation and outflow of CSF*. Such a coupling is also consistent with the work by Williams et al., which showed a coupling between CBV increase and CSF-inflow decrease in response to neuronal activation (Williams et al., 2023). Our findings are also consistent with findings by Zimmermann et al. which showed an increase in CSF outflow that is synchronized with vasodilation (Zimmermann et al., 2025).

#### Dose dependence

The wavelength dependence of the PBM response stems from the behaviours of various chromophores in the brain. In the 808 - 1064 nm range, the main chromophores are CCO and water. As suggested by Saeed et al. (Saeed et al., 2025), wavelengths near 980 nm, 1065 nm or longer than 1200 nm may be a more optimal choice as they are at or near water absorption peaks, and thus they can stimulate the dynamics of MLVs efficiently and lead to the acceleration of CSF circulation.

However, our findings do not show a significant wavelength-only dependence, as seen in **Fig. 9** and **Fig. 10**. Instead, our findings suggest a consistent wavelength-fluence interaction for both tPBM and iPBM. Specifically, at both wavelengths, the intermediate irradiance level, instead of the highest irradiance level, produced the highest CSF response (**Table 4 and Table 5**). This biphasic response pattern is generally consistent with previous findings in cortical neurons (Huang and Hamblin, 2019). Lastly, while there was no significant frequency dependence in the CSF response to iPBM, there was a significant frequency-fluence interaction. Specifically, 10 Hz outperformed 40 Hz but only at an irradiance of 150 mW/cm^2^ as reflected by the aqueduct signal. As the same was not seen in the other CSF ROIs, this observation may be insufficient to prove a consistent frequency dependence in the CSF response to tPBM. To summarize, the interaction between irradiance and tPBM response does not follow the simulated predictions (Van Lankveld et al., 2024), and suggests that the dependence of the PBM effect on irradiance is more nuanced and calls for personalized PBM dosimetry. This in-vivo manifestation of the biphasic irradiance dependence warrants further investigation.

#### Melanin dependence

In accordance with the hypothesis and related simulation results, we found that lighter skin tones are associated with greater CSF responses (**Table 4** and **Table 5**). This is an important experimental validation of our Monte Carlo simulations (Van Lankveld et al., 2024), and implies that in order to target similar CSF-based outcomes in individuals with different skin tones, adjustments may have to be made in the irradiance to compensate for the effects of melanin. This is not a consideration for iPBM or any other form of PBM delivery that does not require penetration of the top layers of the epidermis (Salehpour et al., 2019), and should be considered in PBM protocol design.

### tPBM vs. iPBM

As shown in **Fig. 9** and **Fig. 10**, iPBM produced similar CSF flow increases as tPBM, albeit with only 1/20 the power density. While our current data cannot clarify the mechanisms behind this finding, it is nonetheless consistent with the cribriform plate and the olfactory bulb being on the path of the glymphatic-related CSF outflow (Norwood et al., 2019). The cribriform plate is a porous bone formation whose thickness is only a small fraction that of the frontal skull. Thus, light delivery to the microglia via iPBM should suffer from less attenuation than does tPBM. Moreover, recent research suggests a significant interaction between CSF flow and the olfactory system. CSF drains through the cribriform plate along olfactory nerves, connecting directly to lymphatic vessels in the nasal submucosa (Spera et al., 2023; Zakharov et al., 2004). This so-called olfactory CSF conduit may play a crucial role in clearing metabolites like amyloid-β from the brain, with disruptions potentially contributing to Alzheimer’s disease pathogenesis (Ethell, 2014). The peripheral CSF outflow pathway extends beyond the olfactory system, with evidence of CSF flowing along spinal nerves as well (Bechter and Schmitz, 2014). Understanding these pathways may provide insights into neurodegenerative diseases and offer potential therapeutic targets for enhancing CSF-mediated clearance of harmful metabolites from the brain (Ethell, 2014; Spera et al., 2023). Thus, ultra-low-level PBM through iPBM shows promise that should be further explored.

### Limitations

While this study shows experimental evidence of CSF flow modulation by PBM, which are most likely related to PBM-induced vasodilation, we do not explicitly measure cerebral blood volume, nor do we measure lymphatic pumping. Thus, we do not directly illustrate the former mechanism, nor do we explicitly refute the latter mechanism.

Additionally, fMRI measurements of CSF flow do not constitute quantitative measurements of CSF flow, although our theoretical simulations provide some insight into the relationship between the two. In this context, shorter TRs are typically used to maximize the inflow effect. Nonetheless, it is in theory possible to observe inflow effects in the lowest slice of the fMRI volume even at higher TRs, and our data confirms this. In our future work, we will endeavour to quantify the CSF flow change using specialized techniques such as phase-contrast imaging.

## Conclusion

Our study demonstrates that both tPBM and iPBM can modulate CSF dynamics and potentially enhance glymphatic clearance. Both iPBM and tPBM were able to produce increases in CSF outflow, seen across multiple CSF regions. Light pulsation frequency, irradiance and wavelength as well as melanin (in the case of tPBM) all influenced the extent of CSF flow. Moreover, iPBM may be a more efficient way than tPBM for eliciting a CSF response, due to its delivery proximity to the olfactory system and its lack of melanin dependence. Further research should be done to confirm this finding and to optimize PBM parameters for therapeutic applications.

## Acknowledgments

This work was supported by funding from the Ontario Centre for Innovation, the Natural Sciences and Engineering Research Council of Canada and Vielight Inc.

## Supplementary Materials

**Figure S1.**
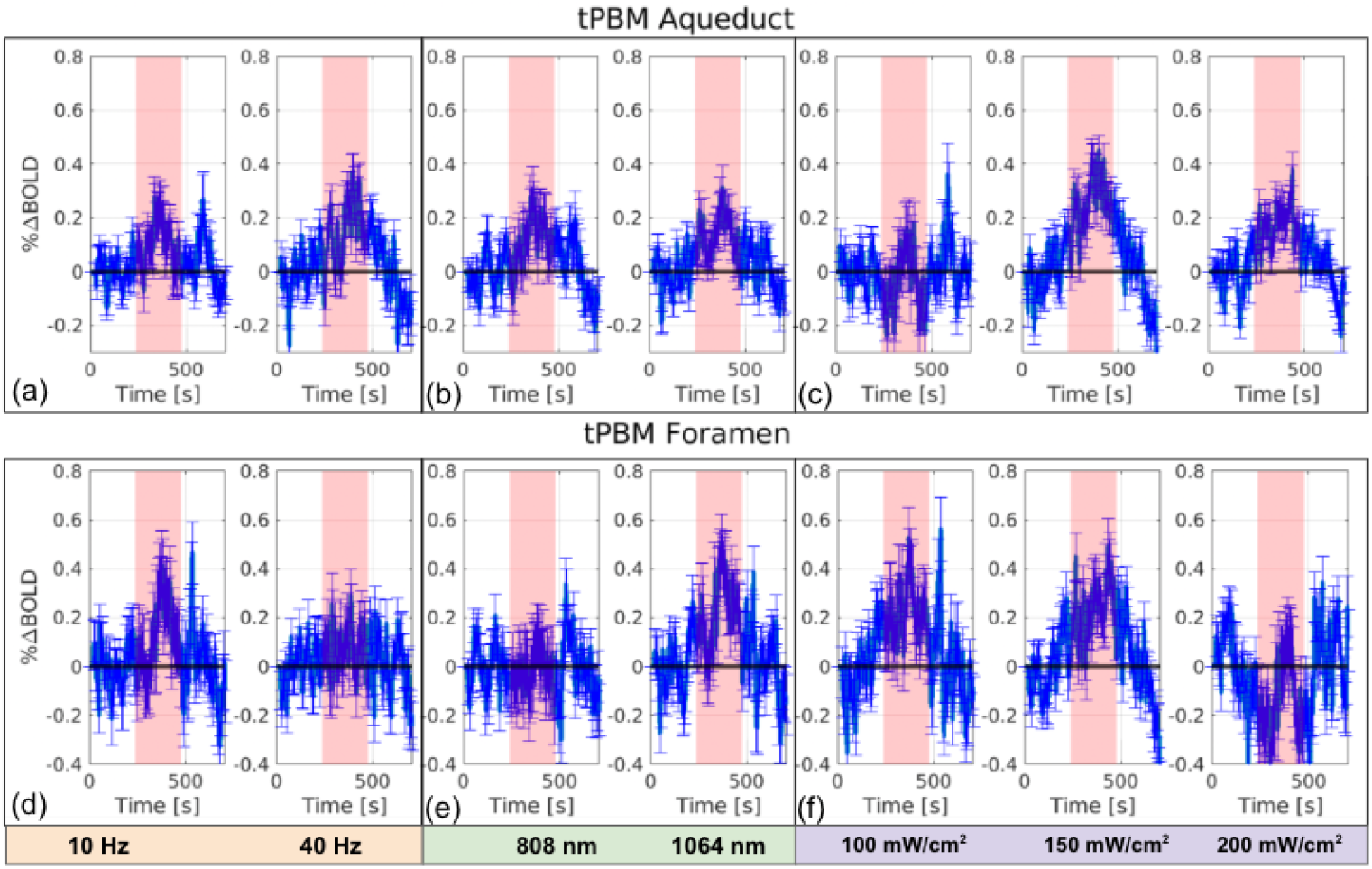
tPBM CSF responses in the cerebral aqueduct and foramen magnum. The Mean BOLD fMRI time courses are shown in blue. In each average, the time courses from all scans of all participants are included, and grouped by the variable of interest only (collapsing across all other variable dimensions). (a) and (d) are grouped by frequency, (b) and (e) grouped by wavelength and (c) and (f) grouped by fluence. The error bars represent the standard error.

**Figure S2.**
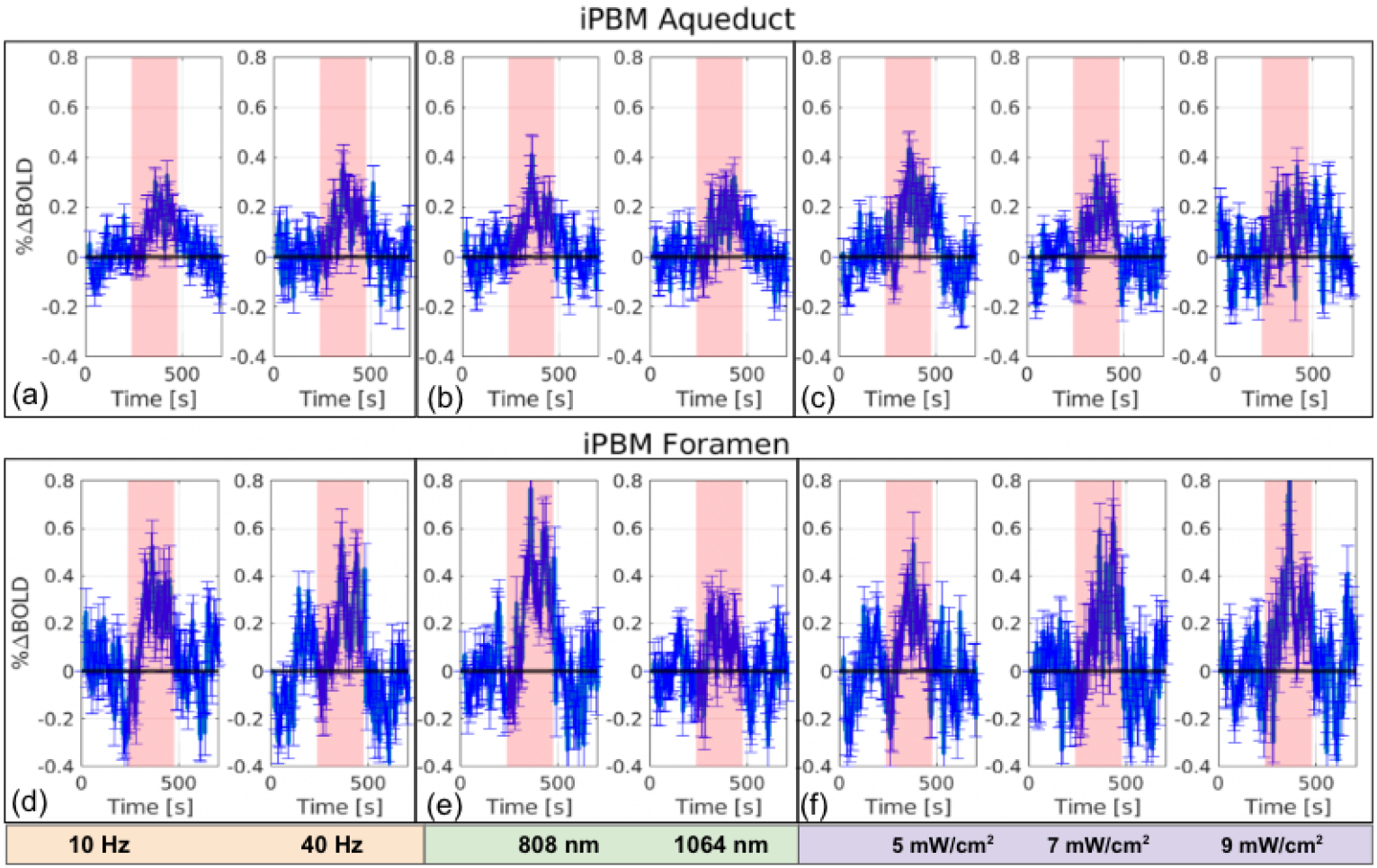
iPBM CSF responses in the cerebral aqueduct and foramen magnum. The Mean BOLD fMRI time courses are shown in blue. In each average, the time courses from all scans of all participants are included, and grouped by the variable of interest only (collapsing across all other variable dimensions). (a) and (d) are grouped by frequency, (b) and (e) grouped by wavelength and (c) and (f) grouped by fluence. The error bars represent the standard error.

## References

Attarpour, A., Ward, J., Chen, J.J., 2021. Vascular origins of low-frequency oscillations in the cerebrospinal fluid signal in resting-state fMRI: Interpretation using photoplethysmography. Hum. Brain Mapp. 10.1002/hbm.25392

Barry, R.L., Smith, S.A., 2019. Measurement of T2* in the human spinal cord at 3T. Magnetic resonance in medicine 82, 743.

Battal, B., Kocaoglu, M., Bulakbasi, N., Husmen, G., Tuba Sanal, H., Tayfun, C., 2011. Cerebrospinal fluid flow imaging by using phase-contrast MR technique. The British Journal of Radiology 84, 758.

Bechter, K., Schmitz, B., 2014. Cerebrospinal fluid outflow along lumbar nerves and possible relevance for pain research: case report and review. Croat. Med. J. 55, 399–404.

Benjamini, Y., Hochberg, Y., 1995. Controlling the false discovery rate: A practical and powerful approach to multiple testing. J. R. Stat. Soc. Series B Stat. Methodol. 57, 289–300.

Cassano, P., Tran, A.P., Katnani, H., Bleier, B.S., Hamblin, M.R., Yuan, Y., Fang, Q., 2019. Selective photobiomodulation for emotion regulation: model-based dosimetry study. Neurophotonics 6, 015004.

Dmochowski, G.M., Shereen, A.D., Berisha, D., Dmochowski, J.P., 2020. Near-Infrared Light Increases Functional Connectivity with a Non-thermal Mechanism. Cereb Cortex Comm 1. 10.1093/texcom/tgaa004

Ethell, D.W., 2014. Disruption of cerebrospinal fluid flow through the olfactory system may contribute to Alzheimer’s disease pathogenesis. J. Alzheimers. Dis. 41, 1021–1030.

Fultz, N.E., Bonmassar, G., Setsompop, K., Stickgold, R.A., Rosen, B.R., Polimeni, J.R., Lewis, L.D., 2019. Coupled electrophysiological, hemodynamic, and cerebrospinal fluid oscillations in human sleep. Science. 10.1126/science.aax5440

Gao, J.H., Liu, H.L., 2012. Inflow effects on functional MRI. Neuroimage 62, 1035–1039.

Halani, S., Kwinta, J.B., Golestani, A.M., Khatamian, Y.B., Chen, J.J., 2015. Comparing cerebrovascular reactivity measured using BOLD and cerebral blood flow MRI: The effect of basal vascular tension on vasodilatory and vasoconstrictive reactivity. Neuroimage 110, 110–123.

Hamblin, M.R., 2018. Mechanisms and mitochondrial redox signaling in photobiomodulation. Photochem. Photobiol. 94, 199–212.

Hamblin, M.R., Demidova, T.N., 2006. Mechanisms of low level light therapy, in: Hamblin, M.R., Waynant, R.W., Anders, J. (Eds.), Mechanisms for Low-Light Therapy. Presented at the Biomedical Optics 2006, SPIE, p. 614001.

Huang, Y., Hamblin, M., 2019. Photobiomodulation on cultured cortical neurons, in: Photobiomodulation in the Brain. pp. 35–47.

Kim, H.B., Baik, K.Y., Choung, P.-H., Chung, J.H., 2017. Pulse frequency dependency of photobiomodulation on the bioenergetic functions of human dental pulp stem cells. Sci. Rep. 7, 15927.

Li, T., Xue, C., Wang, P., Li, Y., Wu, L., 2017. Photon penetration depth in human brain for light stimulation and treatment: A realistic Monte Carlo simulation study. J. Innov. Opt. Health Sci. 10, 1743002.

Mester, E., Szende, B., Gärtner, P., 1968. [The effect of laser beams on the growth of hair in mice]. Radiobiologia, radiotherapia 9.

Mokri, B., 2001. The Monro-Kellie hypothesis: applications in CSF volume depletion. Neurology 56, 1746–1748.

Nairuz, T., Sangwoo-Cho Lee, J.-H., 2024. Photobiomodulation therapy on brain: Pioneering an innovative approach to revolutionize cognitive dynamics. Cells 13, 966.

Norwood, J.N., Zhang, Q., Card, D., Craine, A., Ryan, T.M., Drew, P.J., 2019. Anatomical basis and physiological role of cerebrospinal fluid transport through the murine cribriform plate. Elife 8. 10.7554/eLife.44278

Oner, Z., Kahraman, A.S., Kose, E., Oner, S., Kavakl, A., Cay, M., Ozbag, D., 2017. Quantitative Evaluation of Normal Aqueductal Cerebrospinal Fluid Flow Using Phase-Contrast Cine MRI According to Age and Sex. The Anatomical Record 300, 549–555.

Pastore, D., Greco, M., Passarella, S., 2000. Specific helium-neon laser sensitivity of the purified cytochrome c oxidase. International journal of radiation biology 76. 10.1080/09553000050029020

Poyton, R.O., Ball, K.A., 2011. Therapeutic photobiomodulation: nitric oxide and a novel function of mitochondrial cytochrome c oxidase. Discovery medicine 11.

Ramakrishnan, P., Joshi, A., Fazil, M., Yadav, P., 2024. A comprehensive review on therapeutic potentials of photobiomodulation for neurodegenerative disorders. Life Sci. 336, 122334.

Rasmussen, M.K., Mestre, H., Nedergaard, M., 2022. Fluid transport in the brain. Physiol. Rev. 102, 1025–1151.

Rasmussen, M.K., Mestre, H., Nedergaard, M., 2018. The glymphatic pathway in neurological disorders. Lancet Neurol. 17, 1016–1024.

Saeed, F., Siepker, K.L., Jang, S., Shahdadian, S., Liu, H., 2025. Quantification and stimulation of human glymphatic dynamics:New features of Alzheimer’s disease and effects of brain photobiomodulation. Res. Sq. 10.21203/rs.3.rs-6115809/v1

Salehpour, F., Cassano, P., Rouhi, N., Hamblin, M.R., De Taboada, L., Farajdokht, F., Mahmoudi, J., 2019. Penetration Profiles of Visible and Near-Infrared Lasers and Light-Emitting Diode Light Through the Head Tissues in Animal and Human Species: A Review of Literature. Photobiomodul Photomed Laser Surg 37, 581–595.

Salehpour, F., Khademi, M., Bragin, D.E., DiDuro, J.O., 2022. Photobiomodulation Therapy and the Glymphatic System: Promising Applications for Augmenting the Brain Lymphatic Drainage System. Int. J. Mol. Sci. 23. 10.3390/ijms23062975

Saucedo, C.L., Courtois, E.C., Wade, Z.S., Kelley, M.N., Kheradbin, N., Barrett, D.W., Gonzalez-Lima, F., 2021. Transcranial laser stimulation: Mitochondrial and cerebrovascular effects in younger and older healthy adults. Brain Stimul. 14, 440–449.

Semyachkina-Glushkovskaya, O., Fedosov, I., Zaikin, A., Ageev, V., Ilyukov, E., Myagkov, D., Tuktarov, D., Blokhina, I., Shirokov, A., Terskov, A., Zlatogorskaya, D., Adushkina, V., Evsukova, A., Dubrovsky, A., Tsoy, M., Telnova, V., Manzhaeva, M., Dmitrenko, A., Krupnova, V., Kurths, J., 2024. Technology of the photobiostimulation of the brain’s drainage system during sleep for improvement of learning and memory in male mice. Biomed. Opt. Express 15, 44–58.

Semyachkina-Glushkovskaya, O., Penzel, T., Poluektov, M., Fedosov, I., Tzoy, M., Terskov, A., Blokhina, I., Sidorov, V., Kurths, J., 2023. Phototherapy of Alzheimer’s disease: Photostimulation of brain lymphatics during sleep: A systematic review. Int. J. Mol. Sci. 24, 10946.

Shahdadian, S., Wang, X., Liu, H., 2024. Directed physiological networks in the human prefrontal cortex at rest and post transcranial photobiomodulation. Sci. Rep. 14, 10242.

Spera, I., Cousin, N., Ries, M., Kedracka, A., Castillo, A., Aleandri, S., Vladymyrov, M., Mapunda, J.A., Engelhardt, B., Luciani, P., Detmar, M., Proulx, S.T., 2023. Open pathways for cerebrospinal fluid outflow at the cribriform plate along the olfactory nerves. EBioMedicine 91, 104558.

Tak, S., Wang, D.J.J., Polimeni, J.R., Yan, L., Chen, J.J., 2014. Dynamic and static contributions of the cerebrovasculature to the resting-state BOLD signal. Neuroimage 84, 672–680.

Van Lankveld, H., Chen, J.X., Zhong, X.Z., Chen, J.J., 2025. Consistent and real-time BOLD fMRI response to pulsed transcranial and intranasal photobiomodulation. , in: OHBM 2025 Annual Meeting. Presented at the OHBM 2025 Annual meeting.

Van Lankveld, H., Mai, A.Q., Lim, L., Hosseinkhah, N., Cassano, P., Jean Chen, J., 2024. Simulation-based dosimetry of transcranial and intranasal photobiomodulation of the human brain: the roles of wavelength, power density and skin colour. bioRxiv. 10.1101/2024.04.05.588330

Wang, D.J., Hua, J., Cao, D., Ho, M.-L., 2023. Neurofluids and the glymphatic system: anatomy, physiology, and imaging. Br. J. Radiol. 96, 20230016.

Wang, X., Dmochowski, J.P., Zeng, L., Kallioniemi, E., Husain, M., Gonzalez-Lima, F., Liu, H., 2019. Transcranial photobiomodulation with 1064-nm laser modulates brain electroencephalogram rhythms. Neurophotonics. 10.1117/1.nph.6.2.025013

Wang, X., Tian, F., Reddy, D.D., Nalawade, S.S., Barrett, D.W., Gonzalez-Lima, F., Liu, H., 2017. Up-regulation of cerebral cytochrome-c-oxidase and hemodynamics by transcranial infrared laser stimulation: A broadband near-infrared spectroscopy study. J. Cereb. Blood Flow Metab. 37, 3789–3802.

Wansapura, J.P., Holland, S.K., Dunn, R.S., Ball, W.S., Jr., 1999. NMR relaxation times in the human brain at 3.0 tesla. J. Magn. Reson. Imaging 9, 531–8.

Williams, S.D., Setzer, B., Fultz, N.E., Valdiviezo, Z., Tacugue, N., Diamandis, Z., Lewis, L.D., 2023. Neural activity induced by sensory stimulation can drive large-scale cerebrospinal fluid flow during wakefulness in humans. PLoS Biol. 21, e3002035.

Yan, B., Zhou, J., Yan, F., Gao, M., Tang, J., Huang, L., Luo, Y., 2025. Unlocking the potential of photobiomodulation therapy for brain neurovascular coupling: The biological effects and medical applications. J. Cereb. Blood Flow Metab. 271678×241311695.

Yang, H.-C.S., Inglis, B., Talavage, T.M., Nair, V.V., Yao, J.F., Fitzgerald, B., Schwichtenberg, A.J., Tong, Y., 2022. Coupling between cerebrovascular oscillations and CSF flow fluctuations during wakefulness: An fMRI study. J. Cereb. Blood Flow Metab. 271678×221074639.

Yang, L., Wu, C., Parker, E., Li, Y., Dong, Y., Tucker, L., Brann, D.W., Lin, H.W., Zhang, Q., 2022. Non-invasive photobiomodulation treatment in an Alzheimer Disease-like transgenic rat model. Theranostics 12, 2205–2231.

Yang, Z., Williams, S., Beldzik, E., Anakwe, S., Schimmelpfennig, E., Lewis, L.D., 2024. Attentional failures after sleep deprivation represent moments of cerebrospinal fluid flow. Neuroscience.

Yoo, S.H., 2021. Intranasal photobiomodulation therapy for brain conditions: A review. Med. Lasers 10, 132–137.

Zakharov, A., Papaiconomou, C., Johnston, M., 2004. Lymphatic vessels gain access to cerebrospinal fluid through unique association with olfactory nerves. Lymphat Res Biol 2, 139–146.

Zein, R., Selting, W., Hamblin, M.R., 2018. Review of light parameters and photobiomodulation efficacy: dive into complexity. J. Biomed. Opt. 23, 1–17.

Zhong, X.Z., Chang, C., Jean Chen, J., 2024. Modulation of neurofluid fluctuation frequency by baseline carbon dioxide in awake humans: the role of the autonomic nervous system. Physiology.

Zimmermann, J., Boudriot, C., Eipert, C., Hoffmann, G., Nuttall, R., Neumaier, V., Bonhoeffer, M., Schneider, S., Schmitzer, L., Kufer, J., Kaczmarz, S., Hedderich, D.M., Ranft, A., Golkowski, D., Priller, J., Zimmer, C., Ilg, R., Schneider, G., Preibisch, C., Sorg, C., Zott, B., 2025. Total cerebral blood volume changes drive macroscopic cerebrospinal fluid flux in humans. PLoS Biol. 23, e3003138.

Zinchenko, E., Navolokin, N., Shirokov, A., Khlebtsov, B., Dubrovsky, A., Saranceva, E., Abdurashitov, A., Khorovodov, A., Terskov, A., Mamedova, A., Klimova, M., Agranovich, I., Martinov, D., Tuchin, V., Semyachkina-Glushkovskaya, O., Kurts, J., 2019. Pilot study of transcranial photobiomodulation of lymphatic clearance of beta-amyloid from the mouse brain: breakthrough strategies for non-pharmacologic therapy of Alzheimer’s disease. Biomed Opt Express 10, 4003–4017.

